# Age pigment lipofuscin causes oxidative stress, lysosomal dysfunction, and pyroptotic cell death

**DOI:** 10.1101/2024.03.25.586520

**Authors:** Tim Baldensperger, Tobias Jung, Tom Heinze, Tanja Schwerdtle, Annika Höhn, Tilman Grune

## Abstract

Accumulation of the age pigment lipofuscin represents a ubiquitous hallmark of the aging process. However, our knowledge about cellular effects of lipofuscin accumulation is potentially flawed, because previous research mainly utilized highly artificial methods of lipofuscin generation. In order to address this tremendous problem, we developed a convenient protocol for isolation of authentic lipofuscin from human and equine cardiac tissue in high purity and quantity. Isolated lipofuscin aggregates contained elevated concentrations of proline and metals such as calcium or iron. The material was readily incorporated by fibroblasts and caused cell death at low concentrations (LC_50_ = 5.0 µg/mL) via a pyroptosis-like pathway. Lipofuscin boosted mitochondrial ROS production and caused lysosomal dysfunction by lysosomal membrane permeabilization leading to reduced lysosome quantity and impaired cathepsin D activity. In conclusion, this is the first study utilizing authentic lipofuscin to experimentally validate the concept of the lysosomal-mitochondrial axis theory of aging and cell death.

## INTRODUCTION

Lipofuscin, also known as ceroid^1^ or age pigment^2^, was first described by Hannover in 1843 as a brown pigment in the perikaryon of aged neurons^3^. Lipofuscin can be detected by the measurement of its autofluorescence using either fluorescence or confocal laser scanning microscopy^4^. In addition, histochemical staining for lipids or lipophilic substances using, e.g., Sudan Black or its biotinylated derivative SenTraGor™ are appropriate tools for lipofuscin detection^5^. Lipofuscin is resistant to lysosomal as well as proteasomal degradation and is not exocytosed^6^. Thus, lipofuscin was found to accumulate intracellularly in many different tissues in a time-dependent manner, but especially in non-dividing cells including neurons, skeletal muscle cells, or cardiac myocytes^7–9^. In conclusion, lipofuscin is accepted as an important biomarker of cellular aging.

According to the lysosomal-mitochondrial axis theory of aging, lipofuscin formation is the result of disrupted lysosomal and mitochondrial function. As mitochondrial function declines with age, it leads to increased production of reactive oxygen species (ROS) and damaged proteins. Lysosomes are unable to efficiently process these damaged components and lose their function, which contributes to the further accumulation of dysfunctional mitochondria^10^. In this vicious cycle lipofuscin is formed from oxidized mitochondrial material^11^ and accumulates in lysosomes after macroautophagic uptake^12^. There is a huge variation in lipofuscin composition depending on specific tissues, but in general it is a complex mixture containing oxidized, cross-linked proteins^13^, lipids^14^, and only minor amounts of carbohydrates^15^. Furthermore, incorporation of metals such as iron, copper, manganese, calcium, or zinc up to about two mass percent has been detected^16^.

The redox active metals incorporated in lipofuscin are an important oxidant source in senescent cells^17^. More dramatically, lipofuscin increases susceptibility to oxidative stress by reduced lysosomal function^18^ and inhibition of 20S proteasome^19^. The inhibition of these two major degradation pathways of oxidized macromolecules causes reduced cell viability^17^. Consequently, lipofuscin accumulation is limiting organismal lifespan and plays an important role in age-related pathologies such as macular degeneration^20^, neurodegenerative diseases^21, 22^, and cardiovascular alterations^23, 24^. The first step towards novel therapeutic approaches against these devastating pathologies is elucidation of the underlying molecular mechanisms. Hence, we utilized authentic and matrix-free lipofuscin core particles isolated from cardiac tissue to study cytotoxic effects of lipofuscin aggregates.

## RESULTS

### Lipofuscin isolation and characterization

We developed a protocol for the isolation of authentic lipofuscin from both human and equine cardiac tissue, as demonstrated in **Fig. 1A**. In contrast to previously published methods, we replaced time-consuming ultracentrifugation steps with conventional centrifugation and additional sonication and proteinase K treatment to effectively eliminate impurities. This strategic modification not only streamlined the process but also facilitated the scalability of the isolation procedure to accommodate substantial quantities of heart tissue of up to several hundred grams. Human samples were procured from 4 male and 4 female donors with an average age of 85 years. Between 50 and 330 µg lipofuscin per gram lean heart tissue was extracted from human samples, while only 40 – 60 µg lipofuscin per gram lean heart tissue was extracted from equine hearts. The material isolated from human **(Fig. 1B)** and equine **(Fig. 1C)** cardiac tissue was characterized by confocal microscopy in transmission light mode. Human and equine lipofuscin aggregates appeared under the microscope as round dark granules with diameters ranging from 1 to 5 µm. Further comparison of the confocal micrographs by ImageJ software indicated approximately 25 % higher mean particle size (p = 0.0022) of human compared to equine lipofuscin **(Fig. 1D)**. Human **(Fig. 1E)** as well as equine lipofuscin **(Fig. 1F)** emitted fluorescence in the DAPI channel after excitation by a 405 nm laser. The highest fluorescence intensity was observed for human as well as equine lipofuscin after 365 nm excitation resulting in a broad fluorescence spectrum with a peak of blue light emission between 440 and 500 nm **(Fig. 1G)**.

**Figure 1:**
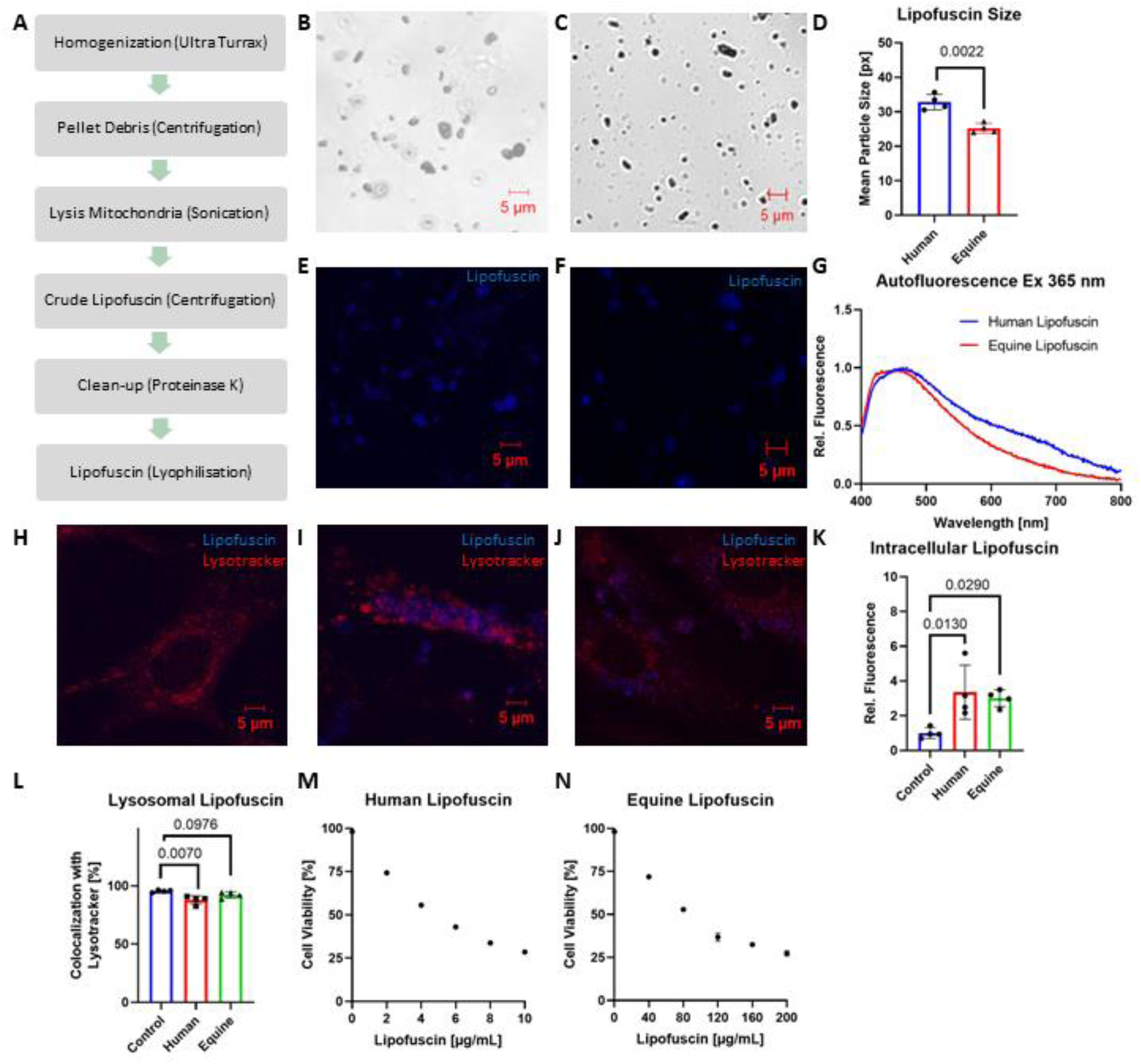
Isolation and characterization of human and equine lipofuscin. **A,** Schematic protocol for isolation of lipofuscin from cardiac tissue. **B,C,** Representative images of human and equine lipofuscin analyzed by confocal microscopy in transmission light channel, respectively. **D,** Comparison of mean lipofuscin particle size by confocal microscopy. Each dot is an individual sample consisting of at least 20 pictures and error bars represent mean ± standard deviation. Statistical analysis was performed by Welch’s t-test. **E, F,** Representative images of human and equine lipofuscin analyzed by confocal microscopy in DAPI channel (excitation 405 nm, emission 410 - 585 nm), respectively. **G,** Fluorescence spectra of human and equine lipofuscin suspended at 1 mg/mL in water after excitation at 365 nm. **H, I, J,** Representative images of FF95 fibroblasts stained by LysoTracker™ Deep Red before and 24 h after exposure to human or equine lipofuscin, respectively. Micrographs were recorded by confocal microscopy using z-stack scans with 1 µm slices through the cells to localize lipofuscin aggregates. Lipofuscin fluorescence was measured between 410 – 585 nm after 405 nm excitation and LysoTracker™ Deep Red was detected between 638 – 756 nm after 633 nm excitation. **K,** Relative quantitation of intracellular lipofuscin by confocal microscopy in FF95 fibroblasts before and after exposure to human or equine lipofuscin. Each dot is an individual sample consisting of at least 20 pictures and error bars represent mean ± standard deviation. Statistical analysis was performed by one-way ANOVA followed by Tukey’s post hoc test. **L,** Colocalization of intracellular lipofuscin with lysosomes. Each dot is an individual sample consisting of at least 20 pictures and error bars represent mean ± standard deviation. Statistical analysis was performed by one-way ANOVA followed by Tukey’s post hoc test. **M**, **N,** Dose-dependent decrease of FF95 fibroblast viability 24 h after exposure to human or equine lipofuscin, respectively. Dead cells with impaired cell membranes were detected by flow cytometry instantly after propidium iodide staining (n = 3).

The toxicological characterization of lipofuscin was conducted within a cellular context utilizing FF95 fibroblasts. These cells had a very low background of blue fluorescence and their lysosomes were stained by LysoTracker™ Deep Red **(Fig. 1H)**. Upon the introduction of human **(Fig. 1I)** or equine **(Fig. 1J)** lipofuscin into the culture medium for 24 h, an incorporation of blue fluorescent particles into the cellular interior was observed through confocal microscopy by employing a z-stack scan methodology with 1 µm sections spanning the cells. Quantitation of intracellular lipofuscin **(Fig. 1K)** by confocal microscopy in the DAPI channel indicated a mean increase of cellular autofluorescence by 100 – 400 % after exposure to human (p = 0.0130) or equine (p = 0.0290) lipofuscin. Lipofuscin fluorescence of untreated cells had a colocalization factor of 95 % with lysosomes **(Fig. 1L)**. After incorporation of human or equine lipofuscin this colocalization dropped to 88 % (p = 0.0070) or 92 % (p = 0.0976), respectively. While morphology, fluorescence spectra, and cellular uptake of both lipofuscin species were comparable, we observed huge differences in toxicity for human **(Fig. 1M)** and equine **(Fig. 1N)** lipofuscin by flow cytometry after propidium iodide staining. Human lipofuscin (LC_50_ 5.0 µg/mL) was about 15 times more toxic in fibroblasts compared to equine lipofuscin (LC_50_ 71.4 µg/mL).

Lipofuscin particles and respective freeze-dried heart tissue underwent hydrolysis under harsh acidic conditions to determine amino acid composition **(Tab. 1)** and metal content **(Tab. 2)** in human and equine samples. In general, no statistically significant differences between both lipofuscin species were detected regarding amino acid composition. The most abundant amino acid in human lipofuscin hydrolysates was glutamic acid with 15 % of total amino acid content and branched-chain amino acids leucine and isoleucine were the least abundant with 1 % each. Proline was significantly (p = 0.0331) enriched in human lipofuscin (14 %) compared to whole heart protein (8 %), while all other amino acids remained unchanged. In terms of metal content, calcium (Ca) and iron (Fe) unequivocally emerged as the dominant constituents within human (5229 mg/kg and 525 mg/kg, respectively) and equine (2741 mg/kg and 1231 mg/kg, respectively) lipofuscin samples **(Tab. 2)**. Trace elements such as zinc (Zn), copper (Cu), nickel (Ni), and manganese (Mn), alongside the toxic metals lead (Pb) and arsenic (As), exhibited comparable concentrations across both lipofuscin types. However, the most striking result from metal analysis was the enormous accumulation of metals in lipofuscin by factors of 2 – 106 compared to respective freeze-dried heart samples, e.g., iron concentration increased by a factor of 3 in human lipofuscin and a factor of 6 in equine lipofuscin.

**Table 1.** Amino acid composition of lipofuscin and heart samples. Values represent mean ± standard deviation. Significant differences (Welch’s t-test, p < 0.05) between lipofuscin and respective heart samples (human n = 8, equine n = 5) are indicated by an asterisk.

| Amino acid | Human lipofuscin<br>[%] | Human heart<br>[%] | Equine lipofuscin<br>[%] | Equine heart<br>[%] |
| --- | --- | --- | --- | --- |
| Alanine | $13 \pm 3$ | $10 \pm 3$ | $15 \pm 3^*$ | $10 \pm 3$ |
| Arginine | $11 \pm 2$ | $8 \pm 2$ | $11 \pm 2^*$ | $8 \pm 2$ |
| Glutamic acid | $15 \pm 2$ | $18 \pm 5$ | $16 \pm 2$ | $18 \pm 5$ |
| Histidine | $12 \pm 1$ | $13 \pm 4$ | $10 \pm 1^*$ | $14 \pm 4$ |
| Isoleucine | $1 \pm 1$ | $2 \pm 1$ | $1 \pm 1$ | $2 \pm 1$ |
| Leucine | $1 \pm 1$ | $1 \pm 1$ | $1 \pm 1$ | $1 \pm 1$ |
| Lysine | $8 \pm 2$ | $12 \pm 4$ | $9 \pm 2^*$ | $14 \pm 4$ |
| Methionine | $2 \pm 1$ | $3 \pm 1$ | $2 \pm 1$ | $2 \pm 1$ |
| Phenylalanine | $7 \pm 1$ | $7 \pm 2$ | $6 \pm 1$ | $7 \pm 2$ |
| Proline | $14 \pm 3^*$ | $8 \pm 2$ | $16 \pm 3^*$ | $8 \pm 2$ |
| Threonine | $7 \pm 1$ | $9 \pm 2$ | $7 \pm 1$ | $8 \pm 2$ |
| Tyrosine | $5 \pm 1$ | $4 \pm 1$ | $3 \pm 1$ | $3 \pm 1$ |
| Valine | $3 \pm 1$ | $4 \pm 1$ | $3 \pm 1$ | $4 \pm 1$ |

**Table 2.** Metal content of lipofuscin and heart samples by ICP-MS. . Values represent mean ± standard deviation. Significant differences (Welch’s t-test, p < 0.05) between lipofuscin and respective heart samples (human n = 8, equine n = 5) are indicated by an asterisk.

| Analyte | Human lipofuscin<br>[mg/kg] | Human heart<br>[mg/kg] | Equine lipofuscin<br>[mg/kg] | Equine heart<br>[mg/kg] |
| --- | --- | --- | --- | --- |
| Ca | 5229 ± 1964* | 640 ± 488 | 2741 ± 625* | 192 ± 8 |
| Fe | 525 ± 244* | 214 ± 44 | 1231 ± 333* | 212 ± 3 |
| Zn | 169 ± 78* | 103 ± 20 | 261 ± 33* | 74 ± 1 |
| Cu | 75 ± 47* | 28 ± 21 | 80 ± 54* | 17 ± 0.2 |
| Ni | 12 ± 9* | 2.7 ± 3.4 | 13 ± 4* | 0.5 ± 0.1 |
| Mn | 2.4 ± 1.0* | 1.4 ± 1.1 | 5.7 ± 1.9* | 1.1 ± 0.1 |
| Pb | 1.7 ± 1.1* | 0.9 ± 1.3 | 3.7 ± 1.8* | 0.2 ± 0.1 |
| As | 0.3 ± 0.1* | 0.01 ± 0.01 | 0.8 ± 0.1* | 0.01 ± 0.01 |

### Mitochondrial function and oxidative stress

Based on the substantial presence of redox-active metals within lipofuscin, our focus centered on investigating its impact on cellular responses to oxidative stress and mitochondrial functionality. Fibroblasts treated for 24 h with 40 µg/mL equine lipofuscin expressed the same amount of mitochondrial marker protein cytochrome c oxidase subunit IV (COX IV) **(Fig. 2A)** compared to untreated control cells (p = 0.7577). MitoTracker™ Green staining **(Fig. 2B)** and subsequent analysis by flow cytometry showed no change in quantity and functionality of mitochondria (p = 0.8737) between groups. However, production of mitochondrial ROS increased significantly (p = 0.0119) by 30 % according to MitoSOX™ Red staining **(Fig. 2C)** after lipofuscin treatment, which is comparable to the rise in ROS levels caused by 40 µM paraquat as a positive control. ROS production after lipofuscin administration induced protein expression of heme oxygenase 1 **(Fig. 2D)** and mitochondrial peroxiredoxin 3 **(Fig. 2E)** by 1.7-fold and 1.3-fold, respectively. These findings underscore the interplay between lipofuscin-induced ROS elevation and, as a consequence, upregulation of key cellular defense mechanisms such as nuclear factor erythroid 2-related factor 2 (NRF2) signaling.

**Figure 2.**
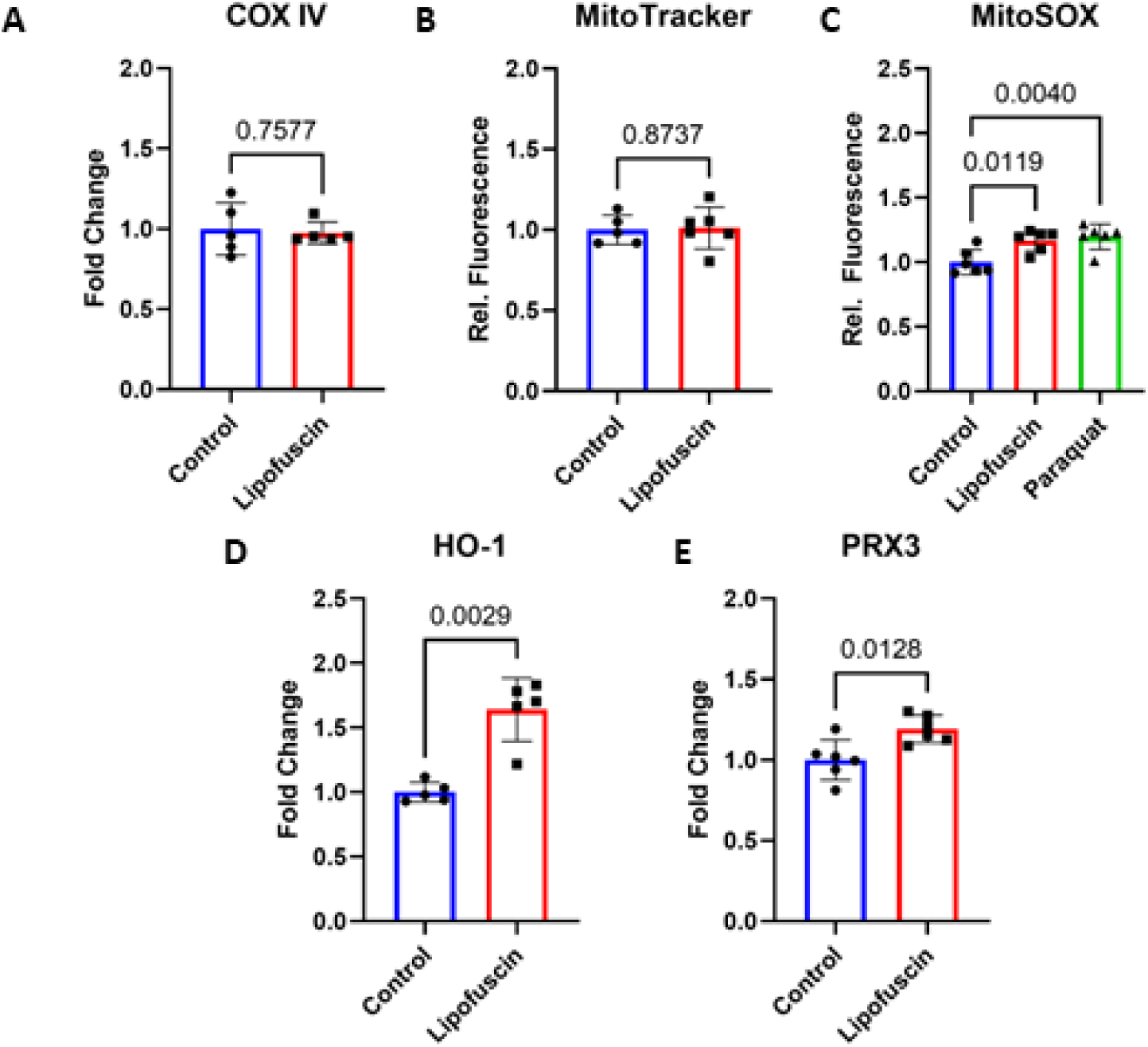
Mitochondrial function and oxidative stress. FF95 fibroblasts treated with 40 µg/mL equine lipofuscin for 24 h were compared to untreated control fibroblasts. Each dot is an individual sample and error bars represent mean ± standard deviation. Statistical analysis was performed by Welch’s t-test (panels A, B, D, E) or one-way ANOVA (panel C) followed by Tukey’s post hoc test. **A,** Protein expression of mitochondrial marker cytochrome c oxidase subunit IV (COX IV) was measured by immunoblotting. **B,** Relative fluorescence of fibroblasts stained by MitoTracker™ Green was analysed by flow cytometry. **C,** Mitochondrial ROS levels were determined by MitoSOX™ Red staining and flow cytometry. A positive control by 40 µM paraquat treatment for 24 h was included. **D, E,** Relative protein expression levels of antioxidative response enzymes heme oxygenase 1 (HO-1) and mitochondrial peroxiredoxin 3 (PRX3) were measured by immunoblots, respectively.

### Lysosomal function

According to the lysosomal-mitochondrial theory of aging impaired lysosomal function due to the accumulation of cellular waste products is a key factor for the escalation of mitochondrial ROS production leading to inflammation and ultimately cell death. Hence, we checked for alterations in lysosomal characteristics, encompassing changes in lysosomal quantity, structural integrity, and the expression and efficacy of vitally important lysosomal enzymes. The total number of lysosomes was significantly reduced by 20 – 30 % as indicated by reduced protein expression (p = 0.0081) of lysosome-associated membrane protein 1 (LAMP1) **(Fig. 3A)** and reduced (p < 0.0001) LysoTracker™ Deep Red fluorescence measured by flow cytometry **(Fig. 3B)**. Moreover, a 2-fold enhancement in the FITC-dextran fluorescence of labeled fibroblast post lipofuscin exposure **(Fig. 3C)** unequivocally indicated the occurrence (p < 0.0001) of lysosomal membrane permeabilization. A similar phenomenon was achieved by 1 mM L-leucyl-L-leucine methyl ester (LLOMe) serving as a positive control. Interestingly, despite decreased lysosomal quantity and the compromised integrity of the lysosomal membrane, the overall intracellular levels of crucial lysosomal enzymes such as lysosomal acid lipase **(Fig. 3D)**, cathepsin B **(Fig. 3E)**, and cathepsin D **(Fig. 3F)** were unaffected by lipofuscin aggregates as confirmed by p-values of 0.2777, 0.5350, and 0.3717, respectively. Perfectly in line with protein expression levels, the enzyme activity of lysosomal acid lipase **(Fig. 3G)** and cathepsin B **(Fig. 3H)** remained constant after lipofuscin treatment. Despite constant protein levels of cathepsin D we detected a tremendous 70 % drop in cathepsin D activity by lipofuscin exposure **(Fig. 3I)**.

**Figure 3.**
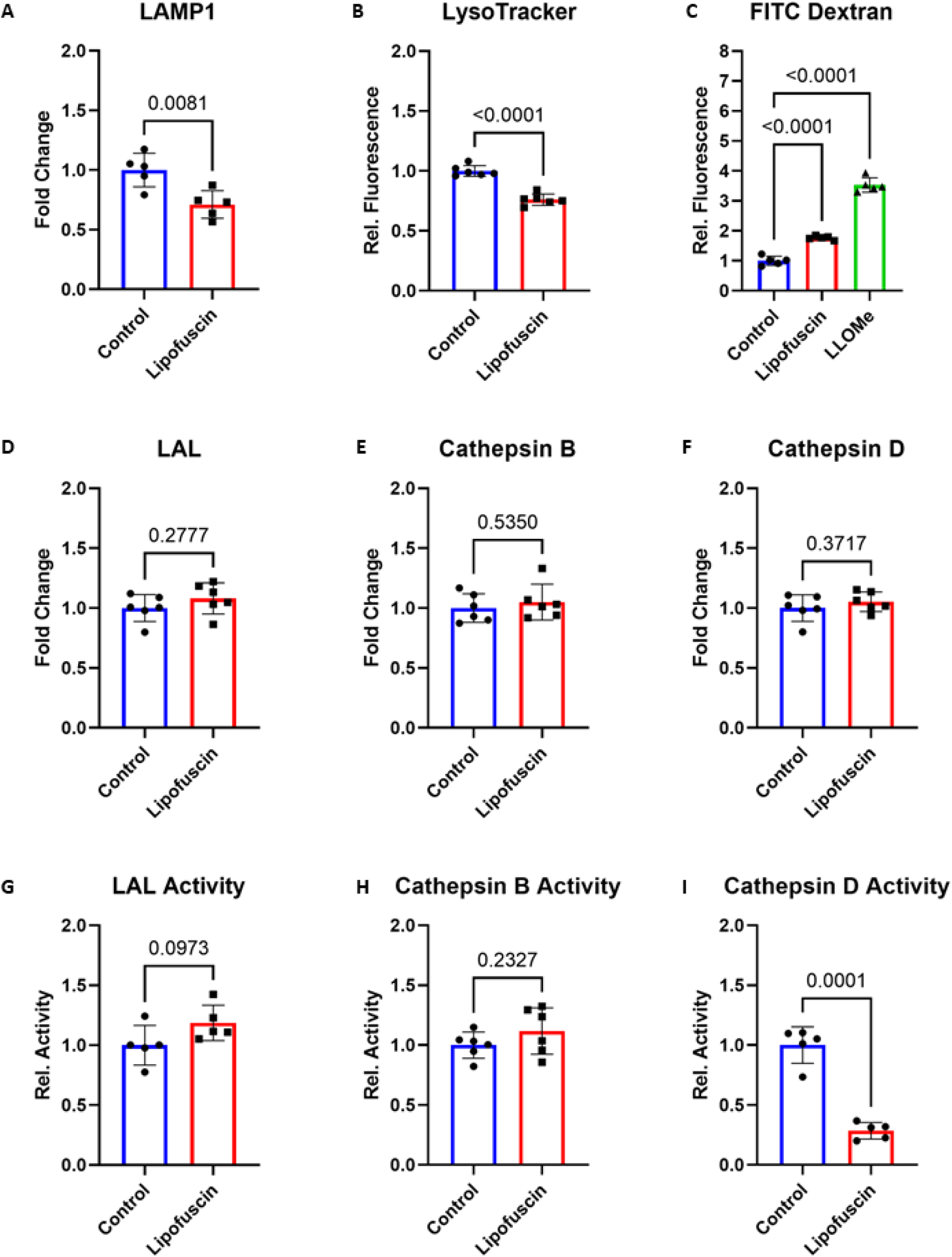
Lysosomal dysfunction. FF95 fibroblasts treated with 40 µg/mL equine lipofuscin for 24 h were compared to untreated control fibroblasts. Each dot is an individual sample and error bars represent mean ± standard deviation. Statistical analysis was performed by Welch’s t-test (panels A, B, D – I) or one-way ANOVA (panel C) followed by Tukey’s post hoc test. **A,** Relative protein concentrations of lysosome-associated membrane protein 1 (LAMP1) were measured by immunoblots. **B,** Relative fluorescence of fibroblasts stained by LysoTracker™ Deep Red was analysed by flow cytometry. **C,** Lysosomal membrane permeabilization was detected by increased fluorescence of FITC-dextran labelled fibroblasts. A positive control by 1 mM L-leucyl-L-leucine methyl ester (LLOMe) treatment for 2 h was included. **D, E, F,** Relative protein expression levels of key lysosomal enzymes lysosomal acid lipase (LAL), cathepsin B, and cathepsin D were measured by immunoblotting, respectively. **G, H, I,** Relative changes in enzymatic activities of key lysosomal enzymes lysosomal acid lipase (LAL), cathepsin B, and cathepsin D were measured in cell lysates using specific fluorosubstrate assays, respectively.

### Cytotoxic mechanism

Last but not least, our attention was directed towards unraveling the toxic mechanism of lipofuscin. Initial investigations encompassed the evaluation of phosphatidylserine exposure on the extracellular cell membrane using FITC-labeled annexin V binding as an marker of apoptotic cell death. The outcome of the annexin V binding assay **(Fig. 4A)** revealed a mere 2 % occurrence of apoptotic fibroblasts in both untreated and lipofuscin-exposed cells. In sharp contrast, exposure to 50 nM staurosporine prompted FITC annexin V binding in 15 % of cells. Immunoblotting of caspase 3 **(Fig. 4B)**, which is the central initiator of apoptosis, detected no change in protein expression between groups (p = 0.2491). Furthermore, exclusively inactive pro-caspase 3 was detected and active forms of cleaved caspase 3 were absent in control and lipofuscin-treated cells. Orthogonal assessment of caspase 3 activation by employing the fluorescent inhibitor of caspase 3 (FLICA) assay supported the previous results and indicated a similar low fraction of fibroblasts (1 – 2 %) with active caspase 3 in control samples and post lipofuscin exposure **(Fig. 4C)**. In contrast, staurosporine activated caspase 3 in 15 % of cells. These results clearly rule out an apoptotic cell death mechanism caused by lipofuscin in fibroblasts. We continued to search for alternative necrotic pathways and utilized immunoblotting to detect a 3.5-fold induction of caspase 1 (p = 0.0014) after lipofuscin exposure **(Fig. 4D)**. A FLICA substrate specific for caspase 1 was used to verify caspase 1 activity **(Fig. 4E)**. Active caspase 1 was nearly absent in control cells, but after lipofuscin treatment 25 % of cells were positive for FLICA caspase 1 (p < 0.0001). Caspase 1 activation marks the initial stage of pyroptotic cell death, a phenomenon recently discovered in macrophages. Consequently, we postulated a pyroptosis-like cell death mechanism for fibroblasts after lipofuscin exposure and analyzed cleavage of gasdermin D by caspase 1. A significant increase (p < 0.0001) of cleaved gasdermin D ranging between 6 – 9 % of total gasdermin D was detected in treated cells **(Fig. 4F)**. In conclusion, our study unveils a previously unknown pyroptosis-like pathway induced by lipofuscin in fibroblasts.

**Figure 4.**
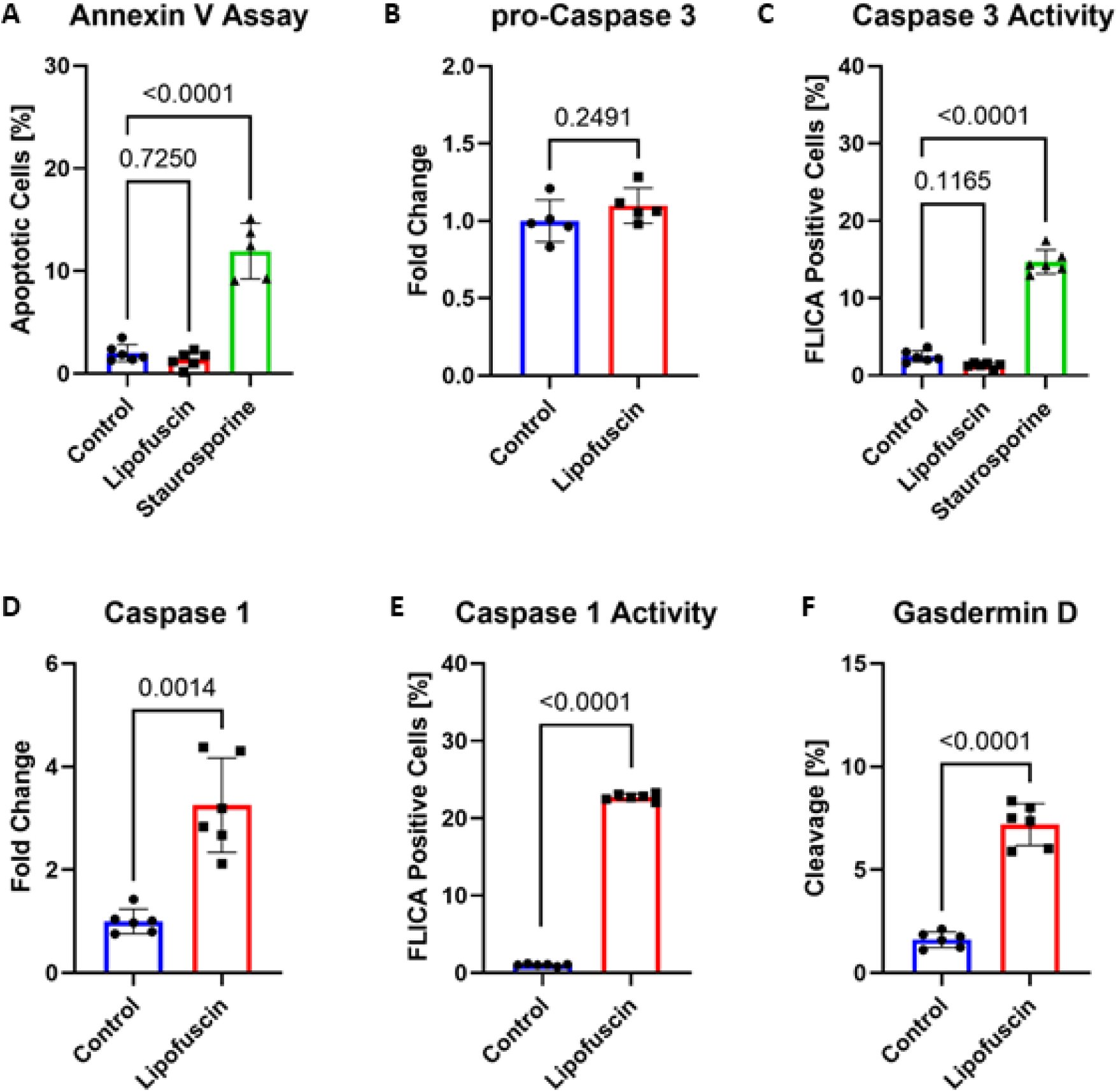
Cytotoxic mechanism. FF95 fibroblasts treated with 40 µg/mL equine lipofuscin for 24 h were compared to untreated control fibroblasts. Each dot is an individual sample and error bars represent mean ± standard deviation. Statistical analysis was performed by one-way ANOVA (panels A and C) followed by Tukey’s post hoc test or Welch’s t-test (panels B and D – F). **A,** Abundance of apoptotic cells was determined by flow cytometry after FITC-annexin V binding assay. A positive control by 50 nM staurosporine treatment for 24 h was included. **B**, Amount of pro-caspase 3 was determined by immunoblotting. **C,** Abundance of apoptotic cells was determined by flow cytometry after fluorescent inhibitor of caspase 3 assay. A positive control by 50 nM staurosporine treatment for 24 h was included. **D,** Relative protein expression levels of caspase 1 were measured by immunoblotting. **E,** Abundance of pyroptotic cells was determined by flow cytometry after fluorescent inhibitor of caspase 1 assay. **F,** Relative share of cleaved gasdermin D was determined by immunoblotting.

## DISCUSSION

The availability of authentic lipofuscin has posed a significant constraint in earlier research efforts. Past studies predominantly employed artificial lipofuscin generated by enhanced oxidative stress (UV irradiation, iron overload, hydrogen peroxide) and inhibited proteolytic pathways (proteasomal and lysosomal small molecule inhibitors) or combination of both approaches in cells and tissue^17, 25^. Authentic lipofuscin derived from tissue remains underutilized due to the limited yield and the labor-intensive traditional isolation methodology involving ultracentrifugation and sucrose density gradients^26, 27^. Hence, we developed a method which is scalable for the isolation of several hundred milligrams of lipofuscin. To ensure purity, heart tissue was chosen as the source material, because to our knowledge no other aggregates than lipofuscin are present contrary to, e.g., brain which contains several different protein aggregates possibly contaminating the isolated material^28^. In order to replace the tedious ultracentrifugation step of traditional isolation protocols we adapted the method described by Hendley et al. utilizing sonication for lysis of mitochondria while preserving lipofuscin granules^29^. However, the original protocol produced preparations with two layers of different colors (light-brown and dark-brown) and a diffuse matrix surrounding the lipofuscin particles under the microscope. Based on the observation that lipofuscin is resistant to proteinase K^30^, we used this enzyme to remove adherent proteins and reduce lipofuscin to its core particle. In our opinion this is a vitally important step, because it is impossible to distinguish whether the surrounding matrix is a natural component of lipofuscin or an artifact from cell lysis and aggregate isolation. The matrix-free core of lipofuscin is the central problem in its accumulation and cellular effects must be elucidated without possibly interfering matrix proteins.

Besides lack of a convenient and scalable method for lipofuscin isolation we solved the problem of limited availability of valuable human tissue by utilizing equine heart as a possible source of lipofuscin. Equine heart is readily available in huge quantities as standard animal feed by several commercial vendors. Horses are mainly used as sport or labor animals and they are slaughtered at a relatively high age leading to relevant quantities of lipofuscin in their hearts. We compared micrographs and fluorescence spectra from human and equine samples isolated by our protocol with cardiac lipofuscin preparations from the literature^29^. Size (1 – 5 µm), morphology (dark-brown and round granules), and fluorescence (excitation 365 nm with broad emission and maximum between 440 – 500 nm) were virtually the same. The chemical analysis of amino acid composition of lipofuscin after acid hydrolysis by LC-MS revealed the same amino acid profile in human and equine heart. Compared to the whole heart, lipofuscin was enriched in proline content, which was noticed by Hendley et al. as well^31^. In contrast to all other proteinogenic amino acids, proline is a secondary amino acid. In combination with protein oxidation and formation of crosslinks the high content of proline possibly explains the extraordinary resistance of lipofuscin aggregates towards proteasomal and lysosomal degradation. Analysis of metal content via ICP-MS identified Ca and Fe as the dominant metals in both lipofuscin species extracted from human and equine heart as well as a comparable trace element profile. Most importantly, we detected a tremendous accumulation of metals in lipofuscin by several magnitudes compared to freeze-dried heart. A similar observation was made by the early work of Jolly et al. in lipofuscin from bovine muscle and brain^16^. In essence, equine heart lipofuscin shares the same morphology, fluorescence, amino acid and metal composition compared to human lipofuscin and we recommend it as a novel model system for in-depth cellular investigations.

Human and equine lipofuscin were internalized to a comparable extent by FF95 fibroblasts as monitored by confocal microscopy utilizing a z-stack scan with 1 µm slices through the cells. A similar phenomenon was reported in literature for uptake of lipofuscin in microglia cells by phagocytosis^32^ and α-synuclein aggregates via endocytosis in THP-1 monocyte cells^33^. In our future research, we intend to comprehensively investigate the precise uptake mechanisms of lipofuscin across diverse cell types. This inquiry is motivated by the potential significance of extracellular lipofuscin resorption from deceased cells, which could play a pivotal role in exacerbating the age-related accumulation of lipofuscin. Human as well as equine lipofuscin addition into the cell culture medium was highly toxic for fibroblasts. Interestingly, human lipofuscin was about 15 times more toxic compared to its equine counterpart. This result was quite unexpected, because both lipofuscin species share very similar properties (morphology, fluorescence spectrum, chemical composition, and cellular uptake) with rather low differences in particle size (25 %) and cytosolic localization (4 %). Beside inter-species variations the higher age of the human donors compared to the horses has to be considered as a possible factor for accumulation of potentially toxic components, which were not detected by our analytical approaches. Despite this small limitation, we consider equine lipofuscin as a superior model system compared to artificial lipofuscin and its high availability opened the opportunity to study cytotoxic mechanisms of authentic lipofuscin in cell culture for the first time.

Based on previous work of our group with artificial lipofuscin^17^ and the high presence of redox-active transition metals in lipofuscin, we expected generation of ROS and activation of anti-oxidative pathways post lipofuscin exposure. Indeed, 30 % elevated ROS levels in mitochondria were detected by MitoSOX™ staining and subsequent flow cytometric analysis of lipofuscin treated cells. A possible explanation for this observation is the import of redox-active metals at the lipofuscin surface into the fibroblasts. The increase of ROS is counteracted by anti-oxidative pathways, e.g., NRF2-signaling and explains the upregulation of mitochondrial peroxiredoxin 3^34^ and heme oxygenase 1^35^.

Another explanation for elevated oxidative stress is lysosomal leakage. The reduction of intracellular LAMP1 and decrease of LysoTracker™ fluorescence were first signs of lysosomal damage after lipofuscin exposure. Lysosomal membrane permeabilization was ultimately proven by enhanced fluorescence of FITC-labeled dextran after addition of lipofuscin to the cell culture medium. This fluorescent dextran normally accumulates in acidic lysosomes. Upon lysosomal rupture the FITC-labeled dextran is leaking into the cytosol. The shift from acidic pH 4.0 in lysosomes to neutral pH 7.4 in cytosol causes a more intense FITC fluorescence and is a widely renowned marker of lysosomal membrane permeabilization^36^. Similar effects on lysosomal integrity were previously described for α-synuclein aggregates^37^. Lysosomal damage had no effects on intracellular levels of lysosomal acid lipase, cathepsin B, and cathepsin D. While activity of lysosomal acid lipase and cathepsin B remained constant, the activity of cathepsin D dropped by about 70 %. This is especially dramatic, because a lack of cathepsin D causes formation of lipofuscin-like autofluorescent granules^38^. Previous studies reported lysosomal membrane permeabilization by iron and leakage of cathepsin D as an inducer of necrotic cell death^39^ and release of cathepsin B into the cytosol as an activator of pyroptosis^40^. Traditionally incited by pathogen-associated molecular patterns (PAMPs), pyroptosis can also be spurred by elevated levels of ROS and lysosomal rupture^41, 42^. Remarkably, our observations regarding the activation of caspase 1 and the concomitant presence of inactive caspase 3 in fibroblasts subjected to lipofuscin treatment in combination with lysosomal rupture and elevated ROS concentrations, closely align with the early phase of pyroptosis. It is noteworthy that these findings diverge from prior studies indicating caspase 3 activation through artificial lipofuscin stimulation^17^, but yet harmonize seamlessly with the distinct form of necrotic cell death induced by N-retinylidene-N-retinyl ethanolamine (A2E), a characteristic fluorophore of lipofuscin extracted from retinal pigment epithelium^43^. Activation of caspase 1 results in cleavage of gasdermin D^44^. The process of gasdermin D cleavage is a hallmark of pyroptotic cell death and generates the gasdermin-N domain. This domain forms 21 nm pores on the cell surface, which initiate membrane rupture and progression of cell death^45, 46^. Hence, we discovered a pyroptosis-like cell death mechanism caused by lipofuscin, which was a previously unknown aspect of lipofuscin toxicity.

In summary, authentic cardiac lipofuscin was used for the first time to elucidate cytotoxicity of lipofuscin. The concerted interplay of oxidative stress induction and lysosomal damage caused by lipofuscin are key factors leading to pyroptotic cell death. The knowledge of these mechanisms is essential to understand lipofuscin toxicity and will strongly facilitate the development of novel therapeutic approaches against cellular loss during aging and age-related diseases.

## METHODS

### Reagents

All chemicals were obtained from Sigma-Aldrich (Taufenkirchen, Germany) and cell culture materials were purchased from Bio & Sell (Feucht, Germany) unless otherwise indicated.

### Lipofuscin isolation

The method described by Hendley et al. was modified for isolation of lipofuscin core particles without matrix proteins^29^. Human heart (donor age 64 – 99 years, mean age 85 years, 4 male and 4 female donors) or equine heart (BARFGOLD, Bassum, Germany) was trimmed free of fat and connective tissue using scissors. The material was further minced by a meat grinder. All preparation steps were performed on ice with pre-cooled buffers unless otherwise stated. About 200 g of minced heart was added to 800 mL suspension buffer (0.75 M sucrose, 0.25 M mannitol, 2 mM TRIS-HCl, 0.2 mM EDTA at pH 7.5) and homogenized by an Ultra-Turrax T25 (IKA, Staufen, Germany) at 24000 RPM for 10 min. Debris was removed by centrifugation in an Avanti JXN-26 with JLA-16.250 rotor (Beckman Coulter, Brea, USA) at 600 RCF for 10 min. The supernatant was filtered by a tea strainer and sonicated by a Sonifier 250 (Branson Ultrasonics, Brookfield, USA) for 5 min. A crude lipofuscin pellet was obtained after centrifugation at 30000 RCF for 10 min. The crude lipofuscin pellet was washed twice by resuspension in wash buffer (0.5 M KCl, 10 mM TRIS at pH 8.0) and pelleted again by centrifugation at 12000 RCF for 10 min. This pellet was resuspended in enzyme buffer (25 M TRIS at pH 8.3) by sonication for 1 min. After addition of 20 mg proteinase K (Carl Roth, Karlsruhe, Germany) the suspension was incubated at 37 °C for 1 h. Finally, pure lipofuscin was pelleted by centrifugation at 12000 RCF for 10 min, washed three times with ultrapure water, and dried by lyophilisation.

### Cell culture

Experiments were performed using FF95 human dermal fibroblasts. Cells were cultivated in Dulbecco’s Modified Eagle Medium (DMEM) supplemented with 10 % fetal calf serum (FCS) and 1 % penicillin/streptomycin at 37 °C in a humidified 5 % CO_2_ atmosphere. Cells were passaged once a week.

### Confocal microscopy

Microscopic verification of lipofuscin was performed by measurement of its autofluorescence by confocal microscopy (Laser scanning microscope 700, Carl Zeiss, Jena, Germany) using the DAPI channel (Excitation: 405 nm; Emission: 498 nm) and 400-fold magnification. FF95 fibroblasts were cultivated in 35 mm glass-bottom dishes (MatTek, Ashland, USA) and stained with LysoTracker™ Deep Red (Thermo Fisher Scientific, Waltham, USA) according to manufacturer protocols to visualize uptake of lipofuscin into cellular lumen. Therefore, a z-stack scan with 12 slices of 1 µm per slice was performed. Lipofuscin particle size, relative concentration, and colocalization with lysosomes were calculated using the ImageJ software package.

### Immunoblot analysis

FF95 fibroblasts were grown in T75 polystyrene flasks (Sarstedt, Nümbrecht, Germany) until 70 – 80 % confluency was reached. Cells were either treated with lipofuscin suspended in PBS (final concentration in cell culture medium 40 µg/mL) for 24 h or the same amount of pure PBS for controls. Fibroblasts were detached by trypsin and washed with PBS. The lysis buffer contained 10 mM Tris-HCl (pH 7.5), 0.9 % Nonidet P-40, 0.1 % SDS and Halt Protease & Phosphatase inhibitor (Thermo Fisher Scientific, Waltham, USA). The protein concentration of the lysate was determined according to the Lowry method. The protein was denatured by reducing 4x Laemmli buffer (0.25 M Tris (pH 6.8), 8 % SDS, 40 % glycerol, 0.03 % SERVA Blue) at 95 °C for 5 min and applied to SDS-PAGE of 12 % (w/v) acrylamide gels followed by electrophoresis and blotting onto 0.45 µm nitrocellulose membrane according to standard procedures. The relative amount of transferred protein per sample was determined after Ponceau S staining by a GS-800 densitometer (Bio-Rad, Hercules, USA) and used as loading control. Immunodetections were performed with the following antibodies at dilutions recommended by the suppliers: rabbit anti-Caspase 1, rabbit anti-Caspase 3, rabbit anti-Gasdermin D, rabbit anti-Lamp 1 rabbit anti-Cathepsin B, rabbit anti-Cathepsin D, rabbit anti-Cox IV, rabbit anti-HO-1, rabbit anti-Peroxiredoxin 3 (Cell Signaling, Boston, USA), rabbit anti-Lysosomal acid lipase, and mouse anti-Gapdh (Abcam, Cambridge, UK). Fluorescent-conjugated secondary antibodies were purchased from Li-Cor Biosciences (Lincoln, USA) and used according to protocols supplied by the manufacturer. The membranes were scanned and stained bands were quantified using an Odyssey® Clx Infrared Imaging System (Li-Cor Biosciences, Lincoln, USA) according to the manufacturer’s instructions.

### Flow cytometry

FF95 fibroblasts were grown in 6-well plates (Sarstedt, Nümbrecht, Germany) until 70 – 80 % confluency was reached. Cells were either treated with lipofuscin suspended in PBS (final concentration in cell culture medium 40 µg/mL) for 24 h or the same amount of pure PBS for controls. DMEM medium was removed and replaced by 1 mL Hanks buffered salt solution per well. For all flow cytometry experiments, cells were stained by Hoechst 33342 (Miltenyi Biotech, Bergisch-Gladbach, Germany) at a final concentration of 15 µg/mL for 15 min. Cell viability was determined by addition of propidium iodide (Miltenyi Biotech, Bergisch-Gladbach, Germany) at a final concentration of 1 µg/mL directly before sample injection. Annexin V-FITC kit (Miltenyi Biotech, Bergisch-Gladbach, Germany) was used according to manufacturer’s protocol to measure apoptosis and a positive control was included by treating FF95 fibroblasts with 50 nM staurosporine. Caspase activity was measured as described in the FAM FLICA caspase 1 and 3 kit (Bio-Rad, Hercules, USA). Lysosomal and mitochondrial content as well as production of ROS were analysed by staining with 50 nM LysoTracker™ Deep Red, 50 nM MitoTracker™ Green, and 5 µM MitoSOX™ Red (Thermo Fisher Scientific, Waltham, USA) for 30 min, respectively. Lysosomal rupture was detected by staining cells for 24 h with 500 kDA FITC-labeled dextran (100 µg/mL) and positive control was 1 mM LLOMe treatment for 2 h as described previously^36^. Stained fibroblasts were washed in autoMACS® Running Buffer (Miltenyi Biotech, Bergisch-Gladbach, Germany) and detached by Trypsin-EDTA at 37 °C for 5 min. Detached cells were centrifuged at 300 RCF for 10 min, resuspended in autoMACS® Running Buffer, and immediately analysed by a MACSQuant® 10 flow cytometer (Miltenyi Biotech, Bergisch-Gladbach, Germany). Nucleated cells were separated from matrix signals by size (forward scatter) and Hoechst 33342 fluorescence. Cell duplets were excluded from the analysis by forward scatter height to area ratio and at least 5000 single cells were counted per experiment.

### Lysosomal activity

FF95 fibroblasts were grown in 6-well plates (Sarstedt, Nümbrecht, Germany) until 70 – 80 % confluency was reached. Cells were either treated with lipofuscin suspended in PBS (final concentration in cell culture medium 40 µg/mL) for 24 h or the same amount of pure PBS for controls. Medium was removed and cells were detached by trypsinization at 37 °C for 5 min. Cells were washed twice in PBS and pelleted at 300 RCF for 10 min. Cells were incubated in 200 μL lysis buffer (1 mM DTT) at 4 °C for 1 h. Following incubation, lysates were sonicated on ice with 10 pulses at 50 % amplitude by UP100 ultrasonic disintegrator (Carl Roth, Karlsruhe, Germany) and subsequently centrifuged at 10000 RCF for 10 min. Protein was quantified by Bradford assay.

For cathepsin activity measurement lysate containing 5 µg of protein was incubated in incubation buffer (150 mM Na-acetate at pH 4.0, 24 mM cysteine, and 3 mM EDTA dihydrate) with specific fluorogenic peptides at 37 °C for 1 h as described previously^47^. Substrates of cysteine protease cathepsin B and aspartyl protease cathepsin D were Z-Phe-Arg-AMC and Mca-Gly-Lys-Pro-Ile-Leu-Phe-Phe-Arg-Leu-Lys(Dnp)-D-Arg-NH_2_ (Enzo Life Sciences, Lörrach, Germany), respectively. The resulting fluorescence (Excitation 360 nm, Emission 460 nm), due to the cleavage and release of fluorophore, was measured in a black 96-well plate by an Infinite M200 (Tecan, Männedorf, Switzerland) for 1 h. The assay was controlled using cathepsin B inhibitor *N*-Acetyl-Leu-Leu-Met and cathepsin D inhibitor pepstatin A.

Activity of lysosomal acid lipase (LAL) was determined in lysate containing 12.5 µg of protein incubated in reaction buffer (0.15 M Na-acetate at pH 4.0, 1 % Triton X-100, and 0.032 % cardiolipin) and release of fluorophore from 4-methylumbelliferyl palmitate. Fluorescence (Excitation 360 nm, Emission 460 nm) was measured in a black 96-well plate by an Infinite M200 (Tecan, Männedorf, Switzerland) at 37 °C for 1 h. The assay was controlled using LAL inhibitor lalistat-2 as described previously^48^.

### LC-MS

Lipofuscin or respective heart protein purified by repetitive 80 % acetone precipitation was hydrolysed in 6 M hydrochloric acid (HCl) containing Supelco stable isotope labeled amino acid mix 1 at 110 °C for 20 h. HCl was evaporated by a Speedvac SPD140DDA vacuum concentrator (Thermo Fisher Scientific, Waltham, USA), samples were reconstituted in MilliQ water, and were filtered through 0.45 μm cellulose acetate Costar SpinX filters (Corning Inc., Corning, USA)^49^.

Analysis of amino acids by LC-MS used the method described by Henning et al.^50^ Separation was performed using an ACQUITY UPLC BEH Amide column (2.1 mm × 100 mm, 1.7 µm) equipped with a VanGuard BEH Amid pre-column (2.1 mm × 5 mm, 1.7 µm) (Waters, Milford USA). Mobile phase A consisted of acetonitrile/MilliQ® water (9:1, v/v) containing 5 mM ammonium formate and 0.3 % formic acid (v/v). Mobile phase B consisted of MilliQ® water containing 25 mM ammonium formate (pH 6.0). Gradient elution started at 98 % A and was held for 4 min. Thereafter, the proportion of solvent B was ramped up to 25 % within 0.5 min. Concentration of 25 % B was maintained between 4.5 and 6.5 min and then solvent B was increased to 40 % within 0.1 min. The 40 % of solvent B was kept constant between 6.6 and 8.0 min. Finally, solvent B was decreased to 2 % within 0.1 min and the column was re-equilibrated with initial conditions for another 1.9 min, resulting in a total runtime of 10 min. The flow rate was constantly set at 0.4 mL/min. Samples were kept in the autosampler at 6 °C and column temperature was set at 35 °C. The mass spectrometer, a Xevo TQ-MS (Waters Corporation, Eschborn, Germany), was operated in ESI-positive mode. Desolvation temperature was set at 600 °C and desolvation gas flow was set at 600 L/h. The source capillary voltage was set at 0.5 kV, cone gas flow was set at 150 L/h and nebulizer gas pressure was set at 7.0 bar.

### ICP-MS

About 25 – 35 mg of lyophilized heart tissue or 1.8 – 2.8 mg of lipofuscin isolated from the corresponding heart samples were weighed into PTFE microwave vessels. Microwave-assisted acid digestion was performed as described before^51^.Therefore, 250 µL of 30 % H_2_O_2_ and 900 µL of 65 % HNO_3_ were used for digestion. 650 µL of deionized H_2_O and 200 µL of an internal standard mix of Rh and Ge, diluted from a 1 g/L stock solution (Rh 100 µg/L; Ge 1000 µg/L, respectively, Carl Roth), and ^77^Se, diluted from a 10 mg/L stock solution (1000 µg/L, ^77^Se 97.20 ± 0.20 %; ^74^Se 0.10 %; ^76^Se 0.40 ± 0.10 %; ^78^Se 2.40 ± 0.10 %; ^80^Se 0.10 %; ^82^Se 0.10 %, certified by Trace Sciences International, Ontario, Canada), were added to the samples. Digestion was performed in a MARS 6 microwave digestion system (CEM, Kamp-Lintfort, Germany) with a ramping phase of 15 min to 200 °C and a holding phase of 20 min. Together with the samples, two replicates of 25 – 34 mg reference material of ERM-BB422 (fish-muscle) and ERM-BB 186 (pig kidney) were digested. After transferring the digested samples into 15 mL tubes, the PTFE vessels were rinsed twice with 1 mL of H_2_O, and the rinse was added to the tubes. Finally, all samples were diluted 1:5 to final concentrations of 2.925 % HNO_3_, 1 µg/L Rh, 10 µg/L Ge, and 10 µg/L ^77^Se. As the ERM-BB reference materials did not report phosphorus concentration, two 1:40 dilutions of an in-house-made KH_2_PO_4_ solution (1.11 g/L) were prepared similar to the reference material. For Mg, P, Ca, Mn, Fe, Co, Ni, Cu, Zn, As, Mo, Cd, and Pb external calibrations were prepared from 1 g/L stock solutions (Carl Roth, Centipur® Merck, or VHG Labs). Se was quantified by isotope dilution analysis (IDA) as described previously^52^. Elements were quantified via inductively coupled plasma-tandem mass spectrometry (ICP-MS/MS, Agilent ICP-QQQ-MS 8800, Agilent Technologies, Waldbronn, Germany) with the following working parameters: 1550 W RF power, plasma gas flow of 15 L/min, make-up gas flow of 0.23 L/min, Ni-cones, MicroMist nebulizer with an Argon flow of 0.99 L/min, Scott-type spray chamber cooled to 2 °C. The following mass-to-charge ratios (m/z) were separated (Q1→Q3) using a He flow of 3 mL/min in the collision reaction cell (CRC) for kinetic energy discrimination (KED): Mg (24→24), Ca (43→43 & 44→44), Mn (55→55), Fe (56→56), Co (59→59), Ni (60→60), Cu (63→63 & 65→65), Zn (64→64 & 66→66), Ge (72→72), Mo (95→95 & 98→89), Rh (103→103), Cd (111→111) and Pb (208→208). In O_2_ mode, with a flow of 30 %, the following m/z were set with a mass-shift of 16, except for Ge and Rh: P (31→47), Ge (72→72), As (75→91), Se (77→93), Se (78→94), Se (80→96), and Rh (103→103). The limit of detection and quantification (LOD and LOQ, respectively) were calculated based on the average concentration of at least 3 technical blanks + 3σ (standard deviation) for LOD or + 10σ for LOQ.

### Statistics

Statistical analysis was performed using Prism 9.0.2 (GraphPad, Boston, USA) software. The data presented in all figures are the mean values ± SD from 5 – 6 replicates. Differences between two groups were assessed by Welch’s t test and p-values of less than 0.05 were considered as significant changes. For multiple comparisons one-way ANOVA was performed and p-values of less than 0.05 were considered as significant changes.

## ACKNOWLEDGEMENTS

The authors sincerely thank Prof. Ingo Bechmann from Institute of Anatomy at University Leipzig for providing access to donated human hearts and those who donated their bodies to science so that our basic research could be performed. Results from such research can potentially increase mankind’s overall knowledge that can then improve patient care. Therefore, these donors and their families deserve our highest gratitude. This work was financially supported by SENS Research Foundation [grant “Lipofuscin Degradation by Bacterial Hydrolases”].

## AUTHOR CONTRIBUTIONS

TB performed experiments, analyzed data and wrote the manuscript. TJ performed confocal microscopy. TH and TS performed elemental analysis by ICP-MS. IB prepared and provided human heart samples. AH and TG developed the concept, supervised the work, and reviewed the manuscript.

## REFERENCES

1. Levine, A. S. et al. Ceroid accumulation in a patient with progressive neurological disease. Pediatrics 42, 583–591 (1968).

2. Samorajski, T., Ordy, J. M. & Keefe, J. R. The Fine Structure of Lipofuscin Age Pkigment in the Nervous System of Aged Mice. J. Cell. Biol. 26, 779–795; 10.1083/jcb.26.3.779 (1965).

3. Hannover, A. Mikroskopiske undersögelser af nervesystemet. Vid Sel naturv. og math (1843).

4. Jung, T., Höhn, A. & Grune, T. Lipofuscin: detection and quantification by microscopic techniques. Methods Mol. Biol. 594, 173–193; 10.1007/978-1-60761-411-1_13 (2010).

5. Evangelou, K. et al. Robust, universal biomarker assay to detect senescent cells in biological specimens. Aging cell 16, 192–197; 10.1111/acel.12545 (2017).

6. Terman, A. & Brunk, U. T. Ceroid/lipofuscin formation in cultured human fibroblasts: the role of oxidative stress and lysosomal proteolysis. Mech. Ageing Dev. 104, 277–291; 10.1016/s0047-6374(98)00073-6 (1998).

7. Beregi, E., Regius, O., Hüttl, T. & Göbl, Z. Age-related changes in the skeletal muscle cells. Z. Gerontol. Geriatr. 21, 83–86 (1988).

8. Brunk, U. & Ericsson, J. L. Electron microscopical studies on rat brain neurons. Localization of acid phosphatase and mode of formation of lipofuscin bodies. J. Ultrastruct. Res. 38, 1–15; 10.1016/S0022-5320(72)90080-9 (1972).

9. Del Roso, A., Tata, V. de, Gori, Z. & Bergamini, E. Transmural differences of lipofuscin pigment accumulation in the left ventricule of rat heart during growth and aging. Aging 3, 19–23; 10.1007/BF03323968 (1991).

10. Terman, A., Gustafsson, B. & Brunk, U. T. The lysosomal-mitochondrial axis theory of postmitotic aging and cell death. Chem.-Biol. Interact. 163, 29–37; 10.1016/j.cbi.2006.04.013 (2006).

11. König, J. et al. Mitochondrial contribution to lipofuscin formation. Redox Biol. 11, 673– 681; 10.1016/j.redox.2017.01.017 (2017).

12. Höhn, A., Sittig, A., Jung, T., Grimm, S. & Grune, T. Lipofuscin is formed independently of macroautophagy and lysosomal activity in stress-induced prematurely senescent human fibroblasts. Free Radic. Biol. Med. 53, 1760–1769; 10.1016/j.freeradbiomed.2012.08.591 (2012).

13. Schutt, F., Bergmann, M., Holz, F. G. & Kopitz, J. Proteins Modified by Malondialdehyde, 4-Hydroxynonenal, or Advanced Glycation End Products in Lipofuscin of Human Retinal Pigment Epithelium. Investig. Ophthalmol. Vis. Sci. 44, 3663–3668; 10.1167/iovs.03-0172 (2003).

14. Bourre, J. M., Haltia, M., Daudu, O., Monge, M. & Baumann, N. Infantile form of so-called neuronal ceroid lipofuscinosis: lipid biochemical studies, fatty acid analysis of cerebroside sulfatides and sphingomyelin, myelin density profile and lipid composition. Eur. Neurol. 18, 312–321; 10.1159/000115095 (1979).

15. Benavides, S. H., Monserrat, A. J., Fariña, S. & Porta, E. A. Sequential histochemical studies of neuronal lipofuscin in human cerebral cortex from the first to the ninth decade of life. Arch. Gerontol. Geriatr. 34; 10.1016/s0167-4943(01)00223-0 (2002).

16. Jolly, R. D., Douglas, B. V., Davey, P. M. & Roiri, J. E. Lipofuscin in bovine muscle and brain: a model for studying age pigment. Gerontology 41 Suppl 2, 283–295; 10.1159/000213750 (1995).

17. Höhn, A., Jung, T., Grimm, S. & Grune, T. Lipofuscin-bound iron is a major intracellular source of oxidants: role in senescent cells. Free Radic. Biol. Med. 48, 1100–1108; 10.1016/j.freeradbiomed.2010.01.030 (2010).

18. Terman, A., Abrahamsson, N. & Brunk, U. T. Ceroid/lipofuscin-loaded human fibroblasts show increased susceptibility to oxidative stress. Experiment. Gerontol. 34, 755–770; 10.1016/s0531-5565(99)00045-5 (1999).

19. Höhn, A. et al. Lipofuscin inhibits the proteasome by binding to surface motifs. Free Radic. Biol. Med. 50, 585–591; 10.1016/j.freeradbiomed.2010.12.011 (2011).

20. Kennedy, C. J., Rakoczy, P. E. & Constable, I. J. Lipofuscin of the retinal pigment epithelium: a review. Eye 9 (Pt 6), 763–771; 10.1038/eye.1995.192 (1995).

21. Braak, E. et al. alpha-synuclein immunopositive Parkinson’s disease-related inclusion bodies in lower brain stem nuclei. Acta Neuropathol. 101; 10.1007/s004010000247 (2001).

22. Cataldo, A. M., Hamilton, D. J. & Nixon, R. A. Lysosomal abnormalities in degenerating neurons link neuronal compromise to senile plaque development in Alzheimer disease. Brain Res. 640, 68–80; 10.1016/0006-8993(94)91858-9 (1994).

23. Waller, B. F. The old-age heart: normal aging changes which can produce or mimic cardiac disease. Clin. Cardiol. 11, 513–517; 10.1002/clc.4960110802 (1988).

24. Kakimoto, Y. et al. Myocardial lipofuscin accumulation in ageing and sudden cardiac death. Sci. Rep. 9, 3304; 10.1038/s41598-019-40250-0 (2019).

25. Gaspar, J., Mathieu, J. & Alvarez, P. 2-Hydroxypropyl-beta-cyclodextrin (HPβCD) reduces age-related lipofuscin accumulation through a cholesterol-associated pathway. Sci. Rep. 7, 2197; 10.1038/s41598-017-02387-8 (2017).

26. Björkerud, S. The isolation of lipofuscin granules from bovine cardiac muscle, with observations on the properties of the isolated granules on the light and electron microscopic levels. J. Ultrastruct. Res. 8, 1–49; 10.1016/S0022--532(063)90001--7 (1963).

27. Ottis, P. et al. Human and rat brain lipofuscin proteome. Proteomics 12, 2445–2454; 10.1002/pmic.201100668 (2012).

28. Davis, A. A., Leyns, C. E. G. & Holtzman, D. M. Intercellular Spread of Protein Aggregates in Neurodegenerative Disease. Annu. Rev. Cell Dev. Biol. 34, 545–568; 10.1146/annurev-cellbio-100617-062636 (2018).

29. Hendley, D. D., Mildavan, A. S., Reporter, M. C. & Strehler, B. L. The properties of isolated human cardiac age pigment. I. Preparation and physical properties. J. Gerontol. 18, 144–150; 10.1093/geronj/18.2.144 (1963).

30. Ng, K.-P. et al. Retinal pigment epithelium lipofuscin proteomics. Mol. Cell. Proteom. 7, 1397–1405; 10.1074/mcp.M700525-MCP200 (2008).

31. Hendley, D. D., Mildavan, A. S., Reporter, M. C. & Strehler, B. L. The properties of isolated human cardiac age pigment. II. Chemical and enzymatic properties. J. Gerontol. 18, 250–259; 10.1093/geronj/18.3.250 (1963).

32. Leclaire, M. D. et al. Lipofuscin-dependent stimulation of microglial cells. Arch. Clin. Exp. Ophthalmol. 257, 931–952; 10.1007/s00417-019-04253-x (2019).

33. Freeman, D. et al. Alpha-synuclein induces lysosomal rupture and cathepsin dependent reactive oxygen species following endocytosis. PloS one 8, e62143; 10.1371/journal.pone.0062143 (2013).

34. Kasai, S., Shimizu, S., Tatara, Y., Mimura, J. & Itoh, K. Regulation of Nrf2 by Mitochondrial Reactive Oxygen Species in Physiology and Pathology. Biomolecules 10; 10.3390/biom10020320 (2020).

35. Loboda, A., Damulewicz, M., Pyza, E., Jozkowicz, A. & Dulak, J. Role of Nrf2/HO-1 system in development, oxidative stress response and diseases: an evolutionarily conserved mechanism. Cell. Mol. Life Sci. 73, 3221–3247; 10.1007/s00018-016-2223-0 (2016).

36. Aits, S., Jäättelä, M. & Nylandsted, J. Methods for the quantification of lysosomal membrane permeabilization: a hallmark of lysosomal cell death. Methods Cell Biol. 126, 261–285; 10.1016/bs.mcb.2014.10.032 (2015).

37. Jiang, P., Gan, M., Yen, S.-H., McLean, P. J. & Dickson, D. W. Impaired endo-lysosomal membrane integrity accelerates the seeding progression of α-synuclein aggregates. Sci. Rep. 7, 7690; 10.1038/s41598-017-08149-w (2017).

38. Escrevente, C., et al. Formation of Lipofuscin-Like Autofluorescent Granules in the Retinal Pigment Epithelium Requires Lysosome Dysfunction. Investig. Ophthalmol. Vis. Sci. 62, 39; 10.1167/iovs.62.9.39 (2021).

39. Lin, Y., Epstein, D. L. & Liton, P. B. Intralysosomal iron induces lysosomal membrane permeabilization and cathepsin D-mediated cell death in trabecular meshwork cells exposed to oxidative stress. Investig. Ophthalmol. Vis. Sci. 51, 6483–6495; 10.1167/iovs.10-5410 (2010).

40. Xie, Z. et al. Cathepsin B in programmed cell death machinery: mechanisms of execution and regulatory pathways. Cell Death Dis. 14, 255; 10.1038/s41419-023-05786-0 (2023).

41. Abais, J. M., Xia, M., Zhang, Y., Boini, K. M. & Li, P.-L. Redox regulation of NLRP3 inflammasomes: ROS as trigger or effector? Antioxid. Redox Signal. 22, 1111–1129; 10.1089/ars.2014.5994 (2015).

42. Chai, R., Li, Y., Shui, L., Ni, L. & Zhang, A. The role of pyroptosis in inflammatory diseases. Front. Cell Dev. Biol. 11, 1173235; 10.3389/fcell.2023.1173235 (2023).

43. Pan, C. et al. Lipofuscin causes atypical necroptosis through lysosomal membrane permeabilization. Proc. Natl. Acad. Sci. USA 118; 10.1073/pnas.2100122118 (2021).

44. Yang, J. et al. Mechanism of gasdermin D recognition by inflammatory caspases and their inhibition by a gasdermin D-derived peptide inhibitor. Proc. Natl. Acad. Sci. USA 115, 6792–6797; 10.1073/pnas.1800562115 (2018).

45. He, W.-t., et al. Gasdermin D is an executor of pyroptosis and required for interleukin-1β secretion. Cell Res. 25, 1285–1298; 10.1038/cr.2015.139 (2015).

46. Santa Cruz Garcia, A. B., Schnur, K. P., Malik, A. B. & Mo, G. C. H. Gasdermin D pores are dynamically regulated by local phosphoinositide circuitry. Nat. Commun. 13, 52; 10.1038/s41467-021-27692-9 (2022).

47. Fernando, R. et al. Age-Related Maintenance of the Autophagy-Lysosomal System Is Dependent on Skeletal Muscle Type. Oxid. Med. Cell. Longev. 2020; 10.1155/2020/4908162 (2020).

48. Hamilton, J., Jones, I., Srivastava, R. & Galloway, P. A new method for the measurement of lysosomal acid lipase in dried blood spots using the inhibitor Lalistat 2. Clin. Chim. Acta 413, 1207–1210; 10.1016/j.cca.2012.03.019 (2012).

49. Baldensperger, T., Jost, T., Zipprich, A. & Glomb, M. A. Novel α-Oxoamide Advanced-Glycation Endproducts within the N6-Carboxymethyl Lysine and N6-Carboxyethyl Lysine Reaction Cascades. J. Agr. Food Chem. 66, 1898–1906; 10.1021/acs.jafc.7b05813 (2018).

50. Henning, T. et al. Pre-Operative Assessment of Micronutrients, Amino Acids, Phospholipids and Oxidative Stress in Bariatric Surgery Candidates. Antioxidants 11, 774; 10.3390/antiox11040774 (2022).

51. Lossow, K. et al. Aging affects sex- and organ-specific trace element profiles in mice. Aging 12, 13762; 10.18632/aging.103572 (2020).

52. Marschall, T. A. et al. Tracing cytotoxic effects of small organic Se species in human liver cells back to total cellular Se and Se metabolites. Metallomics 9, 268–277; 10.1039/c6mt00300a (2017).

